# Automated Spatially Targeted Optical Micro Proteomics (AutoSTOMP) 2.0 identifies proteins enriched within inflammatory lesions in tissue sections and human clinical biopsies

**DOI:** 10.1101/2021.03.19.436192

**Authors:** Bocheng Yin, Laura R Caggiano, Rung-Chi Li, Emily McGowan, Jeffery W Holmes, Sarah E Ewald

**Affiliations:** Department of Microbiology, Immunology and Cancer Biology and the Carter Immunology Center, University of Virginia School of Medicine, Charlottesville VA, USA; Department of Biomedical Engineering, University of Virginia School of Medicine, Charlottesville, VA, USA; Division of Allergy and Clinical Immunology, Department of Medicine, University of Virginia School of Medicine, Charlottesville, VA, USA

## Abstract

Tissue microenvironment properties like blood flow, extracellular matrix or proximity to immune infiltrate are important regulators of cell biology. However, methods to study regional protein expression in context of the native tissue environment are limited. To address this need we have developed a novel approach to visualize, purify and measure proteins in situ using Automated Spatially Targeted Optical Micro Proteomics (AutoSTOMP) 2.0. We previously implemented AutoSTOMP to identify proteins localized to the vacuoles of obligate intracellular microbes at the 1-2 μm scale within infected host cells^1^. Here we report custom codes in SikuliX to specify regions of heterogeneity in a tissue section and then biotin tag and identify proteins belonging to specific cell types or structures within those regions. To enrich biotinylated targets from fixed tissue samples we developed a biochemical protocol compatible with LC-MS. These tools were applied to a) identify inflammatory proteins expressed by CD68^+^ macrophages in rat cardiac infarcts and b) characterize inflammatory proteins enriched in IgG4^+^ lesions in esophageal tissue. These data indicate that AutoSTOMP is a flexible approach to determine regional protein expression in situ on a range of primary tissues and clinical biopsies where current tools are limited.

## INTRODUCTION

Automated spatially targeted optical micro proteomics (AutoSTOMP)^1^ is a proximity-based protein labeling tool that uses standard fluorescence microscopy to visualize structures of interest (SOI). The fluorescence signal is used to identify the pixel coordinates of the SOI and generate a MAP file. The MAP file then guides two-photon excitation of the SOI with UV energy light, which conjugates benzophenone-biotin (BP-biotin) present in the mounting media to any nearby carbon or nitrogen via the benzophenone moiety. Imaging, MAP generation and BP-biotin conjugation is repeated for every field of view and automated using SikuliX icon recognition software. Once biotinylation is complete, unconjugated BP-biotin is washed away. The samples are digested off of the slide. Biotinylated proteins are streptavidin precipitated, then on-bead digested for identification by liquid chromatography mass spectrometry (LC-MS)^1^.

Previously, we demonstrated that AutoSTOMP enriches proteins from the obligate intracellular pathogen *Toxoplasma gondii* (*T. gondii)* within infected human and mouse macrophages^1^. By modifying the MAP to encompass the region surrounding but excluding *T. gondii*, host and *T. gondii* proteins localized to the parasite vacuole membrane were identified. These studies demonstrated that AutoSTOMP can enrich proteins at a 1 μm scale and identify proteins with as little as 1 μg of protein per sample^1^. Proximity-based protein discovery tools that use a label targeting enzyme to biotinylate nearby proteins (BioID, TurboID, APEX) have excellent resolution of ∼10 nm^2,3^. However, they are limited to cell lines or animal models that have tools for genetic modification. Alternatively, in SPPLAT/BAR biotin targeting is mediated by antibodies conjugated to a peroxidase^4,5^. Image guided tagging is a central advantage of AutoSTOMP which allows the user to specify biotin targeting based on co-localization stains or by thresholding, dilating or eroding the boundaries of the image used to guide biotinylation at least ten times the resolution of laser capture micro dissection^6^. Any sample where one or more fluorescent markers (e.g. tags, probes or antibodies) are available to identify SOI is a candidate for AutoSTOMP, so this technique could be a transformative approach to perform localization-dependent protein discovery on a broad range of human clinical specimens.

In practice, however, adapting AutoSTOMP for tissue sections poses several unique challenges compared to cell culture samples. These include: identifying signal versus background in tissues with high or variable autofluorescence, automating the selection of SOI within tissue microdomains rather than elsewhere in a section (particularly for low density SOI), and optimizing the digestion and streptavidin precipitation biochemistry to handle fixed tissues with extensive extracellular matrix networks using detergent compatible with LC-MS^7^. Here we report the AutoSTOMP 2.0 workflow to address the unique demands of in situ proteomics. This includes an updated software analysis package that defines the coordinates of multiple sections on a slide, identifies relevant microdomains of each section and generates a tile array to automate SOI crosslinking. We have developed biochemistry protocols to examine protein enrichment in inflammatory lesions using two disease systems. A rat cardiac infarct model, chosen as a tissue type with extensive extracellular matrix protein crosslinking which poses difficulty for protein purification and human eosinophilic esophagitis (EoE), selected for the small biopsy size and low frequency of lesions in each tissue section.

## MATERIALS AND METHODS

**Note:** Full protocols can be found in “Supporting Methods” of Supporting information.

### Rat cardiac infarct and EoE biopsy collection and staining

Rat myocardial infarcts were induced in 8-week-old male Sprague-Dawley rats (Envigo) by left anterior descending (LAD) coronary artery permanent ligation. 1-week post-surgery the scar region was dissected, frozen in liquid-nitrogen-chilled isopentane, and embedded in OTC. 7 μm cryosections were methanol fixed for 20 minutes on ice and stained with an antibody specific to CD68 (clone: ED1, Bio-Rad). Animal protocols were approved by the University of Virginia Institutional Animal Care and Use Committee.

Six 1 mm biopsies were collected from a patient diagnosed with active EoE (ε15 eosinophils/hpf), according to consensus guidelines, using standard endoscopy procedures^8^. Biopsies were fixed and sectioned as described, and stained with an antibody specific to human lgG4 (clone MRQ-44, Cell Marque). The human study was approved by the University of Virginia Institutional Review Board (IRB), which requires written participant consent (IRB-HSR#19562).

Tissue sections were treated with avidin/biotin blocking kit (SP-2001, Vector Laboratories) then mounted with biotin-dPEG3-benzophenone (BP-biotin, Quanta BioDesign) in 50/50 (v/v) DMSO/water at a concentration of 1 mM. Each slide was prepared immediately prior to AutoSTOMP imaging.

### AutoSTOMP 2.0

Imaging and photo-crosslinking was performed on a LSM880 microscope (Carl Zeiss) equipped with a 25x oil immersion lens (LD LCI Plan-Apochromat 25x/0.8 Imm Korr DIC M27) and a Chameleon multiphoton light source (Coherent). AutoSTOMP 2.0, the upgraded SikuliX (version 1.1.4, http://sikulix.com/) integrated workflow was modified from the previous protocol^1^ and scripted for the tissue to facilitate SOI selection on multiple tissue sections per microscopic slide. Step-by-step instruction and source codes are deposited at https://github.com/boris2008/AutoSTOMP_2.0.git.

Following photo-labeling, each sample was detached from the coverslip. Excess, unconjugated BP-biotin was rinsed with 50/50 (v/v) DMSO/water 3 times, then with water 3 times. The slides were stored at -80 °C before processing replicates in tandem. Rat cardiac sections were lysed in the hydroxylamine lysis buffer^7^ (1 M NH2OH−HCl, 8 M urea, 0.2 M K_2_CO_3_, pH = 9.0) at 45 °C for 17 h to extract proteins from insoluble extracellular matrix. EoE biopsy sections were lysed in DTT/SDS buffer^9,10^ (0.1 M Tris-HCl, 0.1 M DTT, 4% SDS, pH=8.0) at 99 °C for 1 h. Tissue lysates were then diluted 1:10 in TBS-0.1% Tween 20 (TBS-T) then incubated with streptavidin (SA) magnetic beads (Pierce #88817) at room temperature for 1 h. Biotinylated proteins were precipitated by magnet. The unbound proteins were collected as the ‘flow through’ fraction and precipitated with 100% trichloroacetic acid. The biotinylated proteins bound to the beads were washed extensively in TBS reserved for on bead digestion as the AutoSTOMP fraction. The two fractions were desalted and analyzed as previously described using a Thermo Electron Q Exactive HF-X mass spectrometer (MS) at the University of Virginia Biomolecular Analysis Facility^1^.

The raw mass spectra data were parsed by MaxQuant^11^ (versions 1.6.14.0, Max Planck Institute of Biochemistry). The MaxQuant results were then analyzed following the label-free quantification (LFQ) data analysis protocol^11^. Student’s t-test (permutation-based FDR < 0.05) and t-SNE clustering^12^ were applied in Perseus^13^ (versions 1.6.14.0, Max Planck Institute of Biochemistry). The resulting data were plotted in R (www.r-project.org) with the installed packages “ggplot2”, “ggrepel”, “heatmap.2” or using GraphPad Prism (version 8.2.1).

### Immunofluorescence and streptavidin Fluorescence staining

To validate BP-biotin cross-linking, following AutoSTOMP2.0 cross linking some samples were washed and stained with AlexaFluor-594 Streptavidin (#016-580-084, Jackson ImmunoResearch) in TBST. Samples were reimaged to co-localize with the CD68 or lgG4 signal. A full list of antibodies for validation is provided in the Supporting Information.

## RESULTS AND DISCUSSION

### Cardiac infarct macrophages are selectively biotinylated by AutoSTOMP 2.0

To test the ability of the AutoSTOMP protocol to selectively biotinylate structures of interest within tissue sections we first examined a rat myocardial infarction model. In this model, trauma caused by ligation and infiltrating immune cells causes fibroblast activation and deposition of scar tissue that ultimately impairs cardiac function. Macrophage are thought to play a role in the inflammatory regulation and damaged cell turnover in the tissue. One week after surgical ligation on the LAD coronary artery, the infarct region was dissected and cryosectioned for immunofluorescence staining. The infarct region or scar is defined by loss of organized cardiac muscle structure, regions of extracellular matrix and fibroblast expansion as well as infiltrating immune cells. To differentiate between the scar margins, neighboring healthy tissue, low resolution tile scans were performed on adjacent serial sections stained with Hematoxylin and Eosin (H&E) or the macrophage marker CD68 (Figure 1A, yellow border). Using the AutoSTOMP2.0 software module the scar region was tiled into individual fields of view with defined pixel coordinates (Figure 1B). Field of view segmentation and the accuracy of the automated re-imaging program was validated experimentally (Figure S-1).

**Figure 1.**
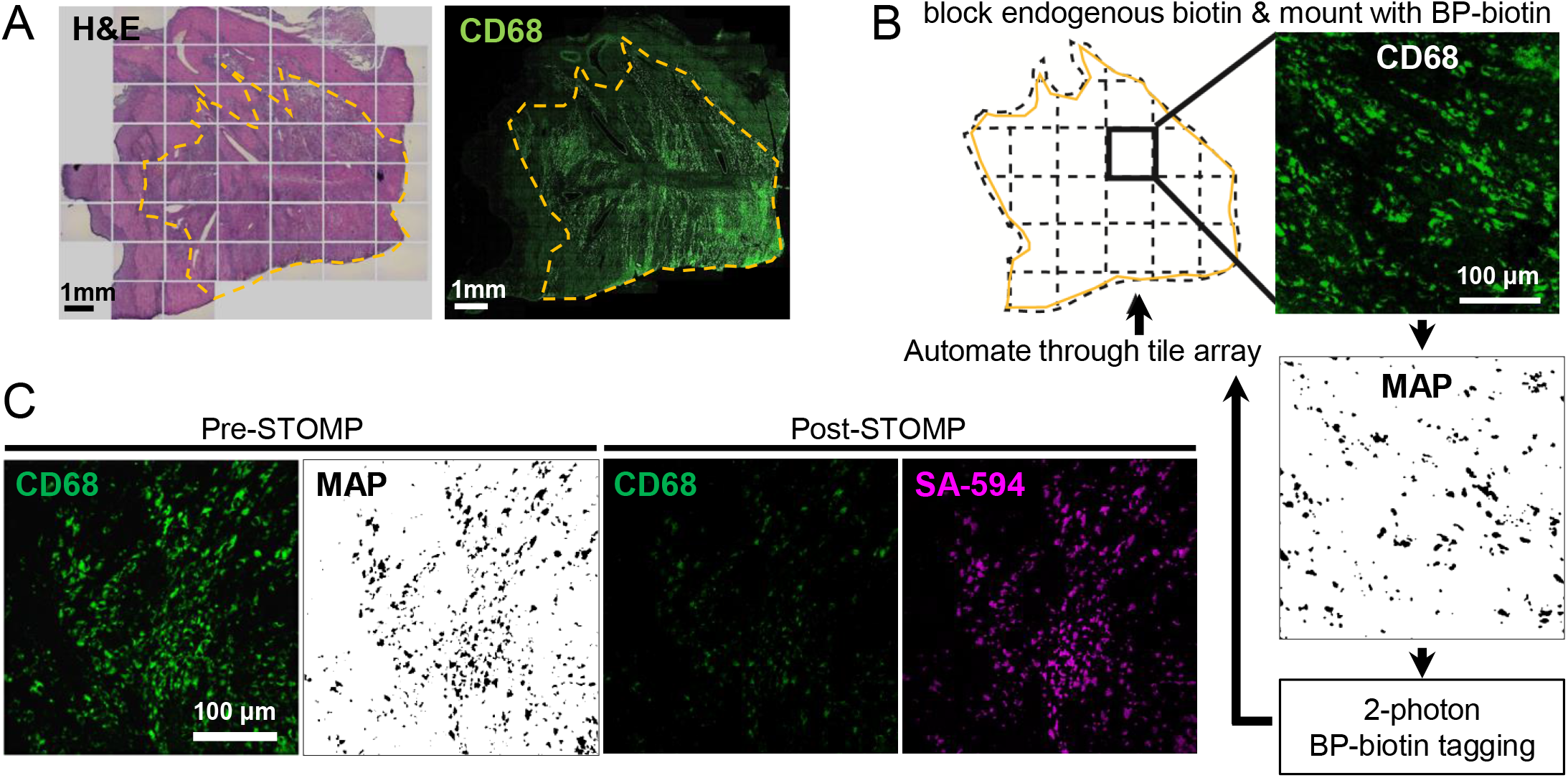
AutoSTOMP identifies rat cardiac infarct borders and biotinylates CD68-associated proteins in tissue sections. **A**, Serial sections of cardiac infarcts were stained with H&E (left) to identify the border of the infarct or the macrophage marker CD68 (green). **B**, The AutoSTOMP 2.0 software divides the infarct region into tiles. In Zeiss Zen Black the CD68 signal is imaged on each tile, exported to FIJI where it is thresholded and used generate a MAP file identifying the pixel coordinates of the CD68+ positive SOI. The MAP file is imported into Zen Black and directs the two-photon to target biotin-BP to the CD68+ SOI. This is automated across the scar. Each field of view measures 340 μm x 340 μm. A typical section contains approximately 550 tiles. **C**, To validate the selective biotinylation of CD68+ SOI (Pre-STOMP) slides were washed, stained with streptavidin-594 then reimaged (Post-STOMP) to assess CD68 (note photobleaching) and streptavidin co-localization.

To biotinylate the CD68^+^ SOI proteins within the scar borders, each tile was imaged at 488 nm (Figure 1B). Each image of the CD68 signal was thresholded and the pixel coordinates were defined in a MAP file. The MAP file then guided the two-photon at 720 nm wavelength, which selectively conjugated BP-biotin to proteins within the CD68^+^ region. This process was fully automated across each field of view in the scar region tile array (Figure 1B). To validate the accuracy of BP-biotin targeting to the CD68^+^ regions, some sections were washed, stained with streptavidin-594 and reimaged (Figure 1C, note photobleaching of Post-STOMP CD68 signal). These data indicate that the AutoSTOMP 2.0 software allows the user to define regions of a tissue section, tile this region into fields of view and accurately image and biotinylate SOI in an automated fashion.

### AutoSTOMP 2.0 enriches macrophage endolysosomal and inflammatory signaling proteins in CD68^+^ regions of cardiac infarcts

To enrich the CD68 SOI proteins, excess unconjugated BP-biotin was washed off of the slide and a sample lysate was prepared. Biotinylated SOI proteins were streptavidin precipitated, eluted by on bead trypsin/LysC digestion and referred to as the ‘CD68’ fraction. To measure protein levels across the entire scar sample, the unbound, ‘flow through’ fraction was also collected (Figure 2A). Of note, the ‘flow through’ fraction is expected to contain CD68^-^ regions as well as any CD68^+^ regions of the section that were deeper than the focal excitation volume of the two-photon (approximately 2.4 μm in the Z axis at 25x magnification^14^). Peptides were identified by LC-MS/MS and Maxquant LFQ^15^ method which measures peak intensity normalized across samples to limit artifacts of run-to-run variability on the LC-MS.

**Figure 2.**
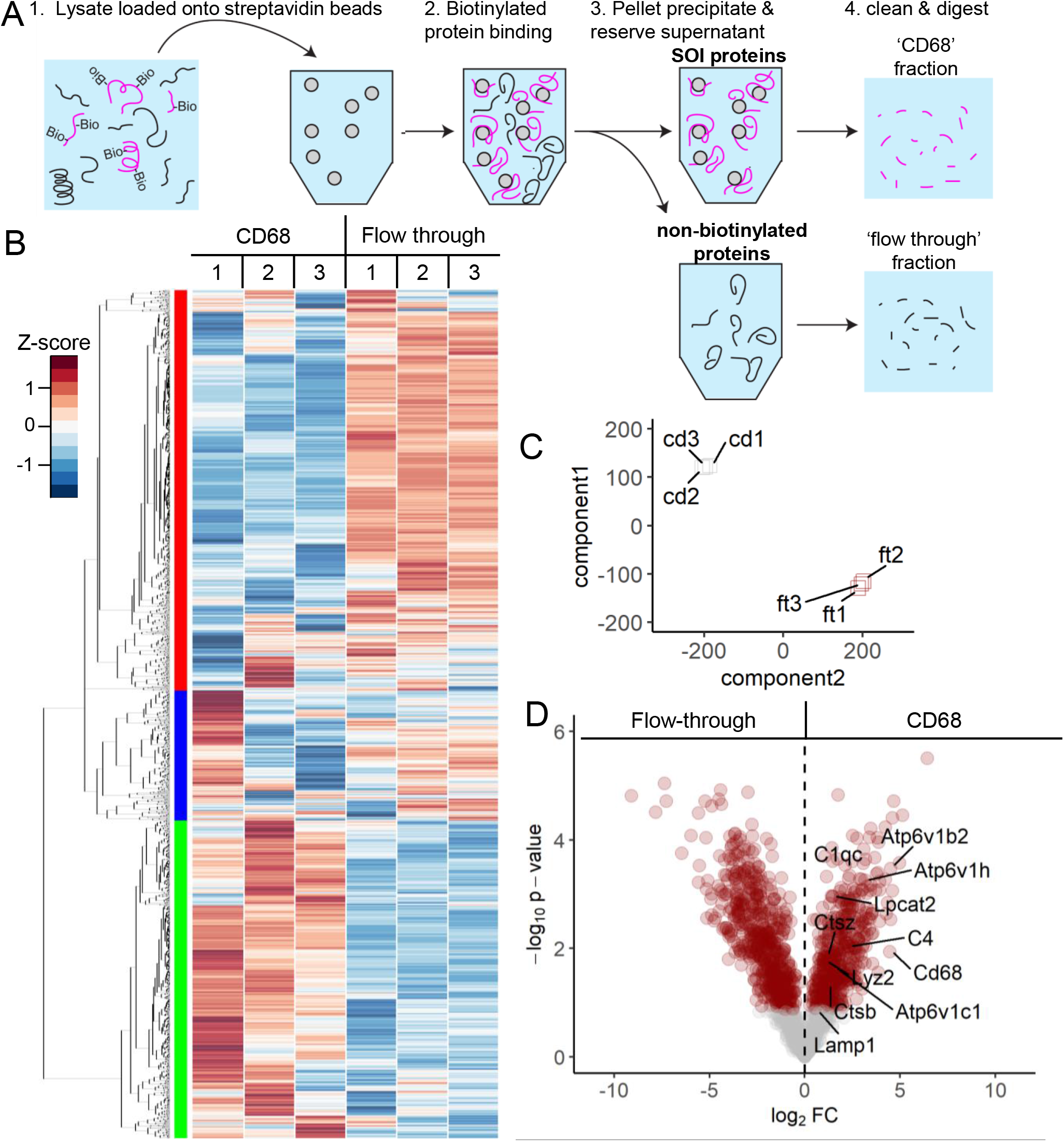
AutoSTOMP selectively enriches macrophage-associated proteins in the CD68^+^ regions of rat cardiac infarcts. CD68^+^ regions of the cardiac infarcts are biotinylated by AutoSTOMP2.0 as described in Figure 1. **A**, After AutoSTOMP 2.0 crosslinking, coverslips are washed of unconjugated BP-biotin and lysed (1). Biotinylated proteins (‘CD68’ fraction) were bound to streptavidin beads (2) and pelleted (3). Unbound proteins are reserved as a ‘flow through’ control (3). Both fractions are trypsin/LysC digested and analyzed by LC-MS(4). **B-D** Protein abundance by MaxQuant label free quantification (MaxLFQ). **B**, Of the 1,671 rat proteins identified, 94.2% were observed with one or more valid readouts in each of the three ‘CD68’ and/or ‘flow through’ fraction replicates. Heat map represents z-score for each protein across all the samples. Hierarchical clustering (left) indicates three enrichment patterns. **C**, T-distributed stochastic neighborhood embedding (t-SNE) analysis of variation among the ‘CD68’ fractions (cd1, cd2, cd3) and ‘flow through’ fractions (ft1, ft2, ft3) belonging to 3 paired replicates. **D**, Of the 1,671 proteins identified, 28.2% of were significantly enriched and 33.7% were significantly lower in the ‘CD68’ fractions relative to the ‘flow through’ (red circles p < 0.05). Plotted as -log_10_ p-value (y axis) versus the log_2_ fold-change (x axis) of the protein abundance averaged between replicates with a false discovery of < 0.1. Macrophage proteins are indicated.

1,671 rat proteins were identified across the ‘CD68’ and ‘flow through’ replicates. Relative expression of each protein across the samples was evaluated by z-score and hierarchical clustering, which indicated that the majority of proteins identified were enriched in ‘CD68’ (Figure 2B, left green bar) or lower in ‘CD68’ fractions (Figure 2B, left red bar) relative to ‘flow through’ fractions. T-distributed stochastic neighborhood embedding (t-SNE)^2^ supported the conclusion that AutoSTOMP 2.0 effectively enriched SOI proteins as there was more similarity within fractions then within each sample (Figure 2C).

Of the 1,671 proteins identified, 28.2% of proteins were more abundant in the ‘CD68’ fraction and 33.7% of proteins were less abundant compared to the ‘flow through’ fractions (Figure 2D, red dots FDR < 0.1). As expected, macrophage markers CD68 (used to guide tagging), Lyz2 (LyzM) and Lpcat2 were enriched in the ‘CD68’ fraction (Figure 2D)^16–18^. The Lyz2 signal co-localized with CD68 (Figure 3A). Consistent with the large phagocytic capacity of macrophages, the lysosomal proteins Lamp1, Acp2, vacuolar ATP-dependent proton pumps (Atp6v1b2, Atp6v1c1, Atp6v1h) and cathepsin (ctsB, ctsz) were enriched in ‘CD68’ fraction (Figure 2D). The CD68^+^ signal partially co-localized with Lamp1 (Figure 3B). The complement proteins C4 and C1q, which are synthesized by macrophages in response to inflammatory stimuli and modulate phagocytic uptake of cargo, were enriched in ‘CD68’ fractions (Figure 2D)^19^. C1qc is expressed by cardiac resident macrophage and co-localized with CD68 (Figure 3C)^20^.

**Figure 3.**
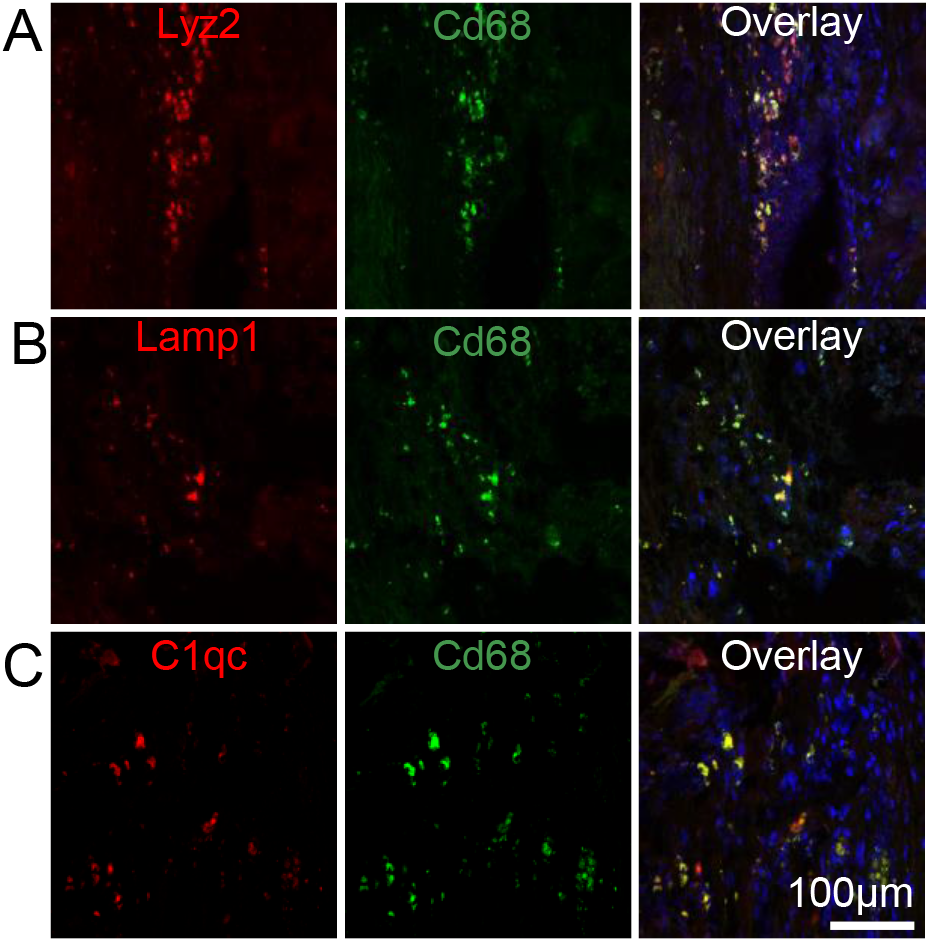
CD68^+^ macrophage partially co-localize with Lyz2, Lamp1 and C1qc in rat cardiac infarcts. 1 week rat cardiac infarcts were stained for CD68 (green) as described in Figure 1 and antibodies specific to the macrophage marker Lyz2 (**A**, red), the lysosomal protein Lamp1 (**B**, red) and the complement protein C1qc (**C**, red). Samples were counter stained with DAPI. N=3.

To identify broader signaling networks associated with CD68^+^ macrophages, the proteins that were significantly different between the ‘CD68’ and ‘flow through’ fractions (Figure 2D, red) were analyzed using David Bioinformatic Resources to annotate Gene Ontology (GO) terms (Figure S-2)^21^. Consistent with inflammatory tissue remodeling, proteins involved in Type 1 interferon cytokine signaling and Glycerol 3-phosphate signaling and metabolism, a pathway that generates lipid signaling mediators of wound healing, were among the most represented ‘biological process’ GO terms in ‘CD68’ fraction (Figure S2A). Proteins regulating amino acid metabolism (aspartate, glutamate), ammonium compound (carnitine), RNA export^22^, and extracellular matrix synthesis were enriched in ‘flow through’ fractions, consistent with muscle regenerative functions of stromal cells (Figure S2B).

Gene set enrichment analysis (GSEA) was also performed on all 1,671 proteins using the REACTOME database (Figure S3)^23,24^. The most highly enriched gene sets in the ‘CD68’ fraction were components of the Eph-ephrin and Fcγ receptor pathways which signal through Rho GTPases (e.g. RhoA, Rac1, and Cdc42) to facilitate actin remodeling and the phagocytic uptake of cargo (Figure S3, red)^25–27^. Interleukin 12 (IL-12), the main macrophage product necessary for IFN-γ expression was also enriched in ‘CD68’ fractions (Figure S3, red)^28,29^. Similar to the results of the GO search, the ‘flow through’ fraction was enriched for proteins belonging to mitochondrial biogenesis and respiration, muscle contraction and extracellular matrix regulation (Figure S3, blue). In summary, these data show that AutoSTOMP2.0 is an effective tool to enrich and measure macrophage proteins in infarcted cardiac tissue sections.

### AutoSTOMP 2.0 enriches granulocyte proteins, eicosanoid inflammatory mediators and glycolytic metabolism machinery from IgG4^+^ inflammatory lesions in esophageal biopsies

We next asked if AutoSTOMP 2.0 could selectively enrich proteins associated with discrete regions of human tissue biopsies. Eosinophilic esophagitis (EoE) is a disease driven by dietary allergens that leads to focal inflammatory lesions within the esophagus, which are characterized by infiltration of eosinophils and mast cells, and increased levels of Th2 cytokines^30^. IgG4 has recently been identified in the esophageal tissue and is increasingly recognized as a relevant feature of this disease^29,30^. However, progress towards understanding disease pathogenesis has been hindered by a lack of well-established animal models, the extremely limited access to samples from the primary site inflammation and heterogeneity in the biopsy tissue^31^. To determine if AutoSTOMP was amenable study EoE pathology, six 1 mm esophagus biopsies were isolated by endoscopy from a patient diagnosed with active EoE (Figure 4-5f). Immunofluorescence staining for lgG4 showed that the lesions measured between 50-300 μm in diameter or approximately 10% of each section (Figure 4A). Using AutoSTOMP 2.0, the boundary of all 6-8 sections per slide was defined by the user to facilitate a low-resolution tile scan (Figure S4A-C). To avoid the high background fluorescence signal from the apical epithelium, each lgG4^+^ SOI was selected by the user (Figure S4D, Figure 4B ‘threshold’ signal vs. ‘MAP’). A tile array of the SOI pixel coordinates (Figure S4E) was then generated to automate BP-biotin tagging (Figure S4F-H). To validate BP-biotin targeting, some slides were reserved for streptavidin-594 staining and reimaged (Figure S5).

**Figure 4.**
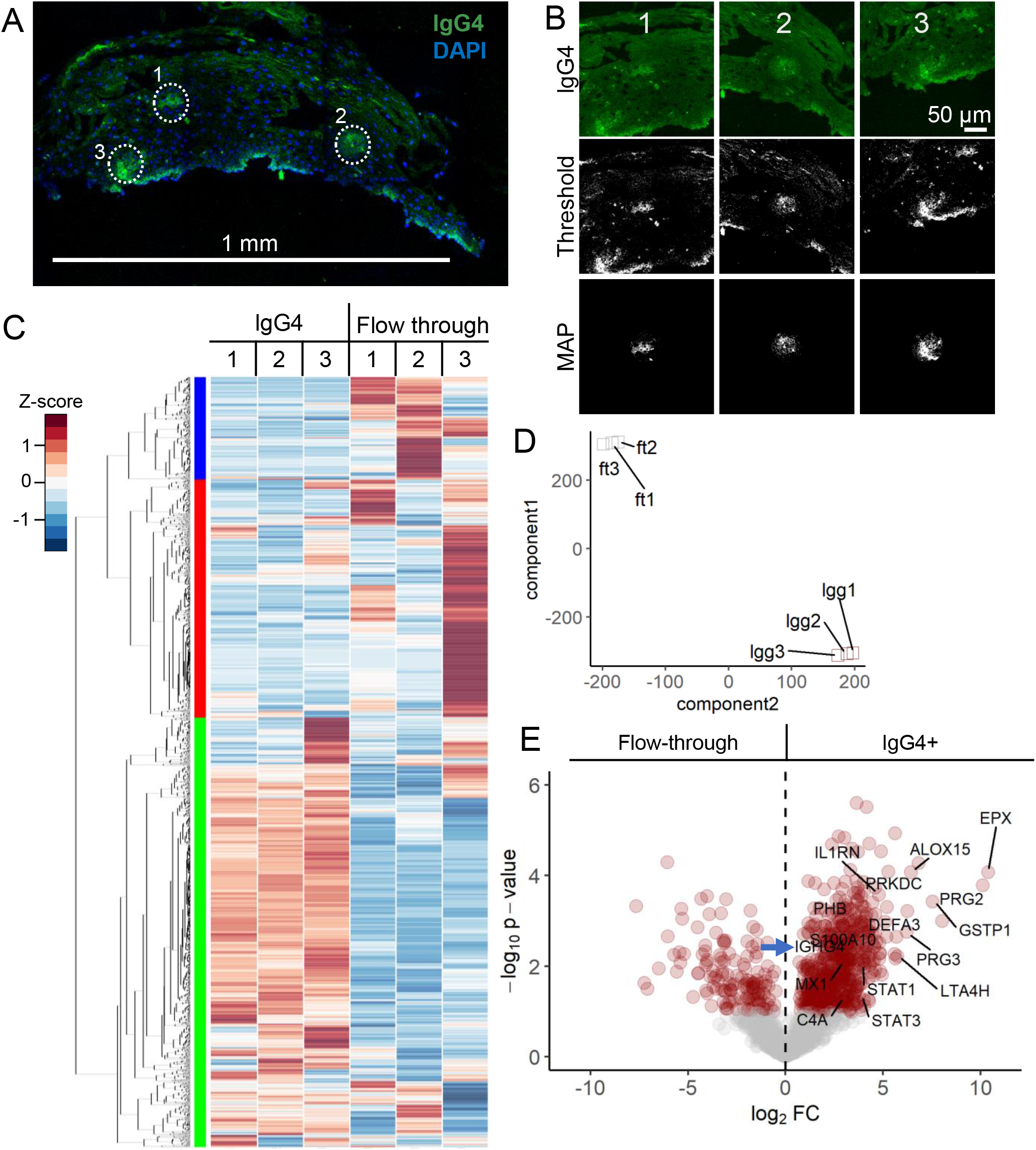
AutoSTOMP identifies eosinophil granule proteins associated with IgG4^+^ lesions in EoE patient esophagus biopsies. **A**, Representative section of an EoE patient biopsy stained with an antibody specific to lgG4 (green) and counterstained with DAPI (blue). IgG4^+^ SOI are indicated by white dots, scale bar = 1 mm. **B**, Three representative SOI tile images, thresholded images and MAP files used to guide biotinylation of each SOI. **C-D**, Biotinylated proteins, the ‘lgG4’ fraction, or the unlabeled ‘flow through’ were isolated and identified as described in Figure 2A. Each sample represents sections pooled from two biopsies. **C**, 2,007 human proteins were identified and plotted by row z-score normalized across all the samples. Hierarchical clustering (left) indicates three enrichment groups. **D**, T-distributed stochastic neighborhood embedding (t-SNE) account for variation among the ‘lgG4’ fractions (lgg1, lgg2, lgg3) and ‘flow through’ fractions (ft1, ft2, ft3) across the three samples. **E**, Of the 2,007 proteins identified, 27.9% of were significantly enriched and 12.3% of the proteins were significantly lower in the ‘lgG4’ fractions relative to the ‘flow through’ fractions (red circles, p < 0.05, FDR < 0.1). Plotted as -log_10_ p-value (y axis) versus the log_2_ fold-change (x axis) of the protein abundance averaged between replicates with a false discovery of < 0.1. IgG4 (IGHG4, blue arrow indicated) is noted with granulocyte proteins (DEFA3, EPX, PRG2, PRG3), inflammatory proteins (IL1RN(aka. IL-1RA), STAT1, STAT3, MX1, C4A, etc.) and lipoxin/prostaglandin proteins (GSTP1, ALOX15, LTA4H).

**Figure 5.**
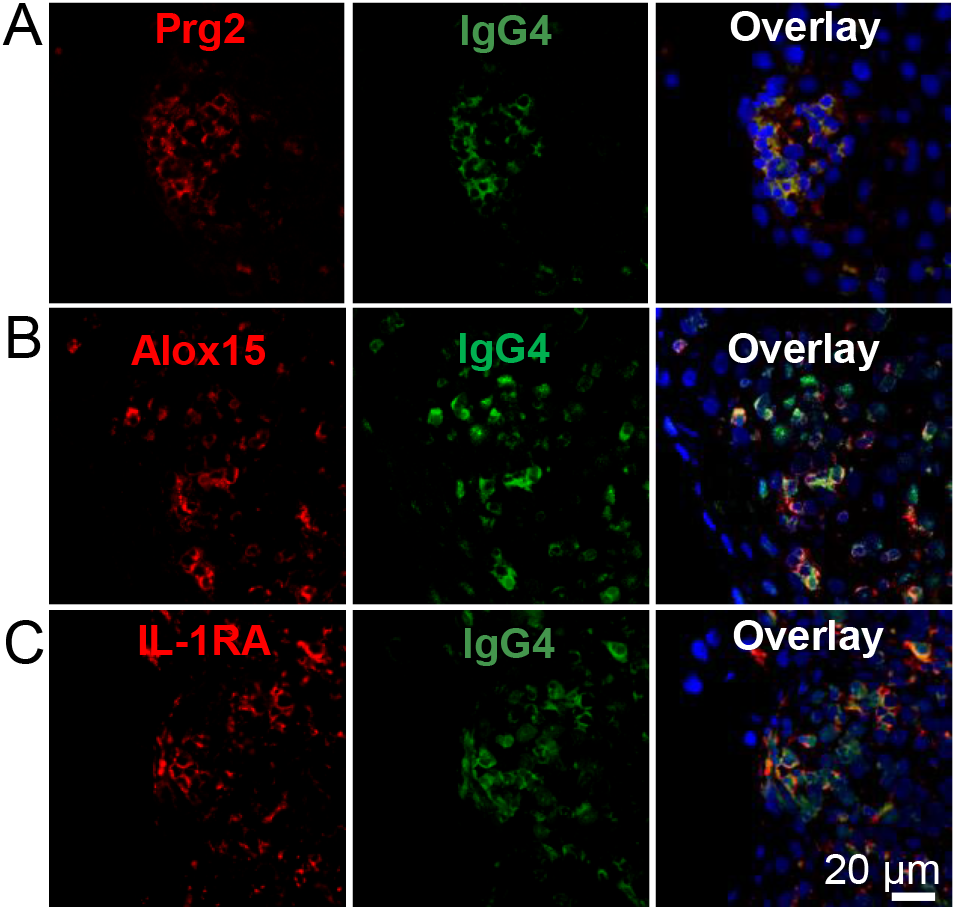
IgG4+ EoE lesions partially co-localize with inflammatory markers. EoE patient biopsy sections were stained for IgG4 (green) as described in Figure 4 and antibodies specific to the proteins Prg2 (**A**, red), Alox15 (**B**, red) and IL-1RA (**C**, red). Samples were counter stained with DAPI. N=3.

To identify the biotinylated proteins enriched in the lgG4^+^ lesions (Figure 4A-B), the sample lysate was incubated with streptavidin beads (‘lgG4’ fraction) and the unbound, ‘flow through’ fraction was reserved as a total protein control as described in Figure 2A. In total, 2,007 human proteins were identified across 3 samples where each sample (1-3) represents paired ‘lgG4’ or ‘flow through’ fraction pooled from 2 biopsies. (Figure 4C). When the level of each protein was evaluated across all samples hierarchical clustering revealed three major groups enriched in ‘IgG4’ fractions (Figure 4C, left green bar) or enriched in ‘flow through’ fractions (Figure 4C, left red & blue bars). Variability in the ‘flow through’ samples accounted for the bimodal clustering of the red and blue groups. This indicated that the lgG4^+^ regions were more similar in protein identity than the stromal cells from each biopsy pair and underscored the selective enrichment of the AutoSTOMP 2.0 procedure. The selectivity of AutoSTOMP was also evaluated by t-SNE, which showed that despite the variability within ‘flow through’ fractions, there was more similarity within fractions than within each sample (Figure 4D).

Of the 2,007 proteins identified in the patient’s biopsies, 27.9% of were significantly enriched in the ‘lgG4’ fraction and 12.3% of the proteins were depleted in ‘lgG4’ fraction relative to the ‘flow through’ (Figure 4E). Granulocyte secretory proteins were among the most highly enriched proteins in the ‘lgG4’ fractions including the proteoglycans Prg2 and Prg3, Defensin DefA3 and eosinophil peroxidase EPX (Figure 4E).^32^ Enzymes regulating the synthesis of eicosanoid lipid inflammatory regulators were also enriched in the ‘IgG4’ fraction including arachidonate-15 lipoxygenase (Alox15), Leukotriene A-4 hydrolase (LTA4H) and glutathione S-transferase P (GSTP1) which modifies prostaglandin A2 upstream of eicosanoid synthesis^33–35^. Mediators of inflammatory cytokine production were also enriched in the IgG4 regions including the transcription factors STAT1 and STAT3, the interferon induced effector MX1, complement protein C4A, and IL-1R antagonist (IL-1RA) consistent with local activation of the inflammatory response. There was a partial overlap in the staining of Prg2, Alox15 and IL-1RA with the IgG4 lesions, confirming their presence in the patient’s EoE lesions as determined by LC-MS (Figure 5).

To identify signaling networks associated with the IgG4+ lesions we searched the significantly differentially expressed proteins (Figure 4E, red) against the ‘biological process’ GO term library. Most of the enriched pathways were characterized by the abundance of proteasome components detected in the IgG4 lesions (Figure S-6A, asterisks). Independent of this proteasome signature, glycolytic metabolic pathways were the most highly represented GO-terms in the ‘IgG4’ fraction, consistent with metabolic demands needed to drive an inflammatory response (Figure S-6, red)^36^. By contrast, the ‘flow through’ fraction was enriched in proteins regulating epithelial turnover and differentiation (Figure S-6, blue)^37^. Although future studies with an expanded number of patients will be needed to draw definitive conclusions about the mediators of EoE, these data indicate that AutoSTOMP 2.0 is an effective protein discovery tool to identify immune effectors from discrete inflammatory foci in human tissue biopsies.

## CONCLUSION

Here we show AutoSTOMP 2.0 facilitates the discovery of regional proteomes in primary tissue sections and human biopsies. In theory, image guided tagging means that almost any structure that can be visualized is a potential target for AutoSTOMP. In practice, however, heterogeneity in SOI size, shape and frequency were barriers to performing STOMP in tissues. The icon recognition software SikuliX overcomes these limitations by allowing the user to automate image capture, image processing and tiling in the discrete software packages. In addition, SikuliX facilitates the experimental timeline, since AutoSTOMP-mediated biotinylation can take several days per replicate. Finally, developing a protocol to analyze protein levels in the flow through has maximized the use of each sample to assess protein enrichment. This will be a critical parameter for future studies with human samples, to evaluate patient-to-patient variability and where sample access is limited.

## ASSOCIATED CONTENT

“Supporting Methods” contains complete AutoSTOMP protocols, methodology and reagent lists. Supplemental Figures (Figure S1-6)

## Author Contributions

B.Y., and S.E.E. designed the experiments. L.C and J.H performed rat cardiac infarcts. R.L. and E.M. collected clinical specimens. B.Y. performed AutoSTOMP experiments and data analysis. B.Y. and S.E.E. prepared the manuscript.

## ACKNOWLEDGEMENTS

We thank Dr. Nicholas E. Sherman and Dr. Jeong-Jin Park for LC-MS services at the W.M. Keck Biomedical Mass Spectrometry Laboratory. This work was supported by NIH R21AI156153, R35GM138381 (SEE), AMH/Allen Institute (JH and SEE) and start-up funds from the University of Virginia SOM and the Emily Couric Cancer Center.

