## supplemental for "Automated Spatially Targeted Optical Micro Proteomics (AutoSTOMP) 2.0 identifies proteins enriched within inflammatory lesions in tissue sections and human clinical biopsies"

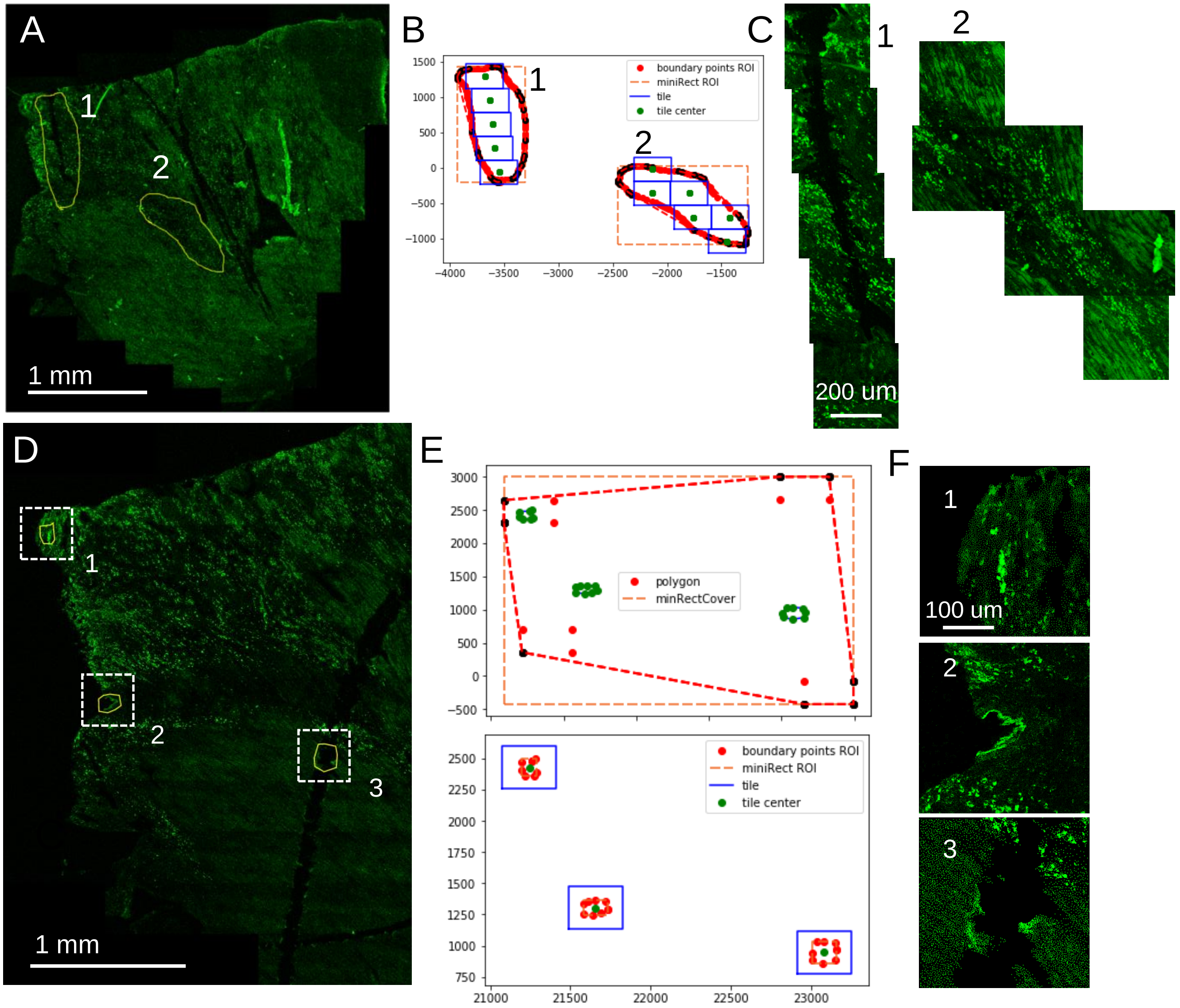


**Figure S1**. **AutoSTOMP 2.0 accurately defines the coordinates of individual fields of view within user defined borders and targets them for re-imaging**. Rat cardiac infarct sections are stained for CD68 as described in Figure 1 and a low resolution tile scan (single tile size, 340 µm x 340 µm) is generated with 25x magnification objective lens. **A-C,** The border of large target regions (greater than 1 field of view) are defined by the user (**A,** yellow line). **B,** The coordinates of each target region are defined (orange line) and broken into individual tiles (blue boxes with green central coordinate). **C,** The coordinates are used to reimage the SOIs and stitched to reproduce the target regions in B. **D-F** The border of small target regions (less than 1 field of view) are defined by the user. (**D,** yellow line) and the pixel coordinates of the perimeter region defined (**E**, top, red dotted line). The center (green) of each tile (purple) is imputed for reimaging the chosen SOIs (red). (**F**) Automated re-imaging of each SOI field of view.

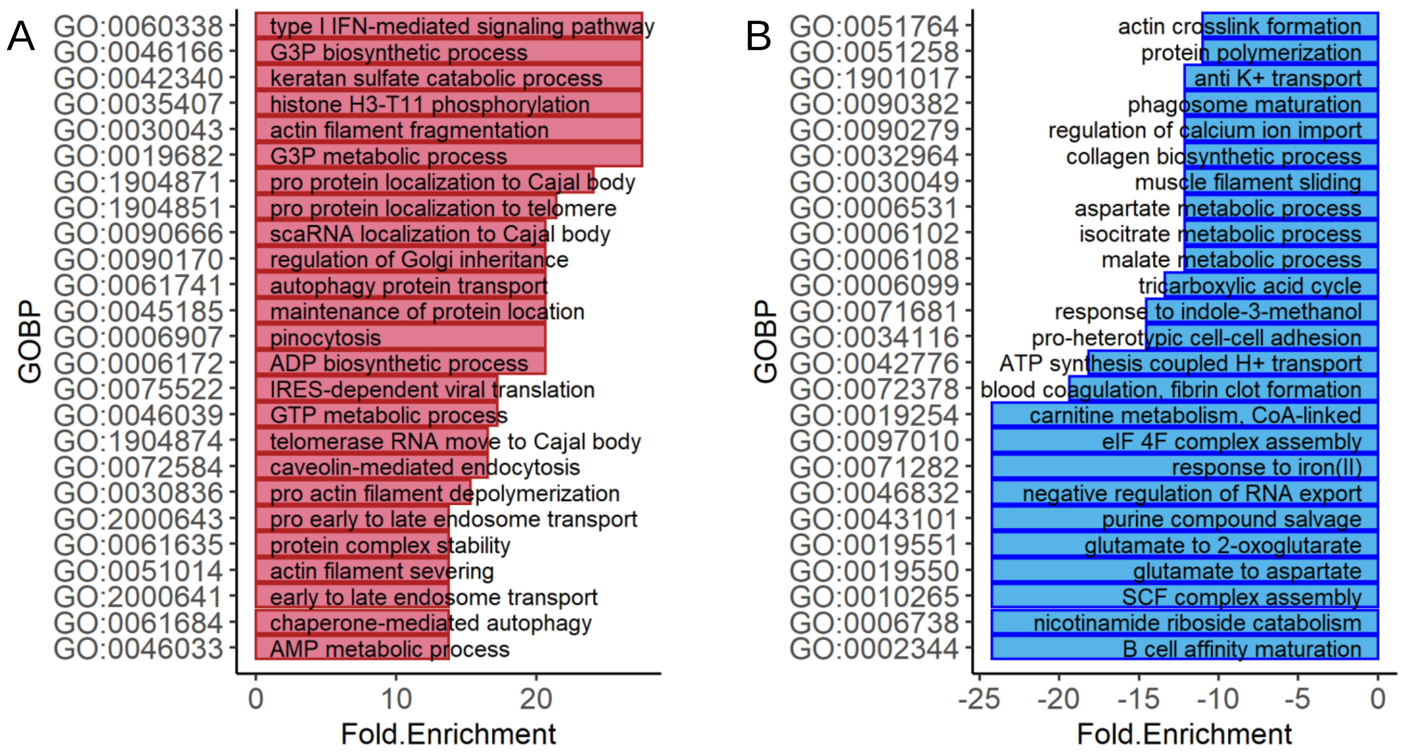


**Figure S2**. **Inflammation regulatory pathways are enriched in ‘CD68’ fractions isolated by autoSTOMP and amino acid metabolism pathways are enriched in ‘flow through’ fractions of rat cardiac infarcts**. Proteins that were significantly enriched in ‘CD68’ fractions (**A**) or the ‘flow through’ fractions (**B**) (Figure 2A, red) were searched against the Gene Ontology (GO) terms Biological Process (BP) database using DAVID Bioinformatics Resource. The top 25 significantly enriched GO terms are displayed in an order ranked by fold enrichment and false discovery rate (FDR).

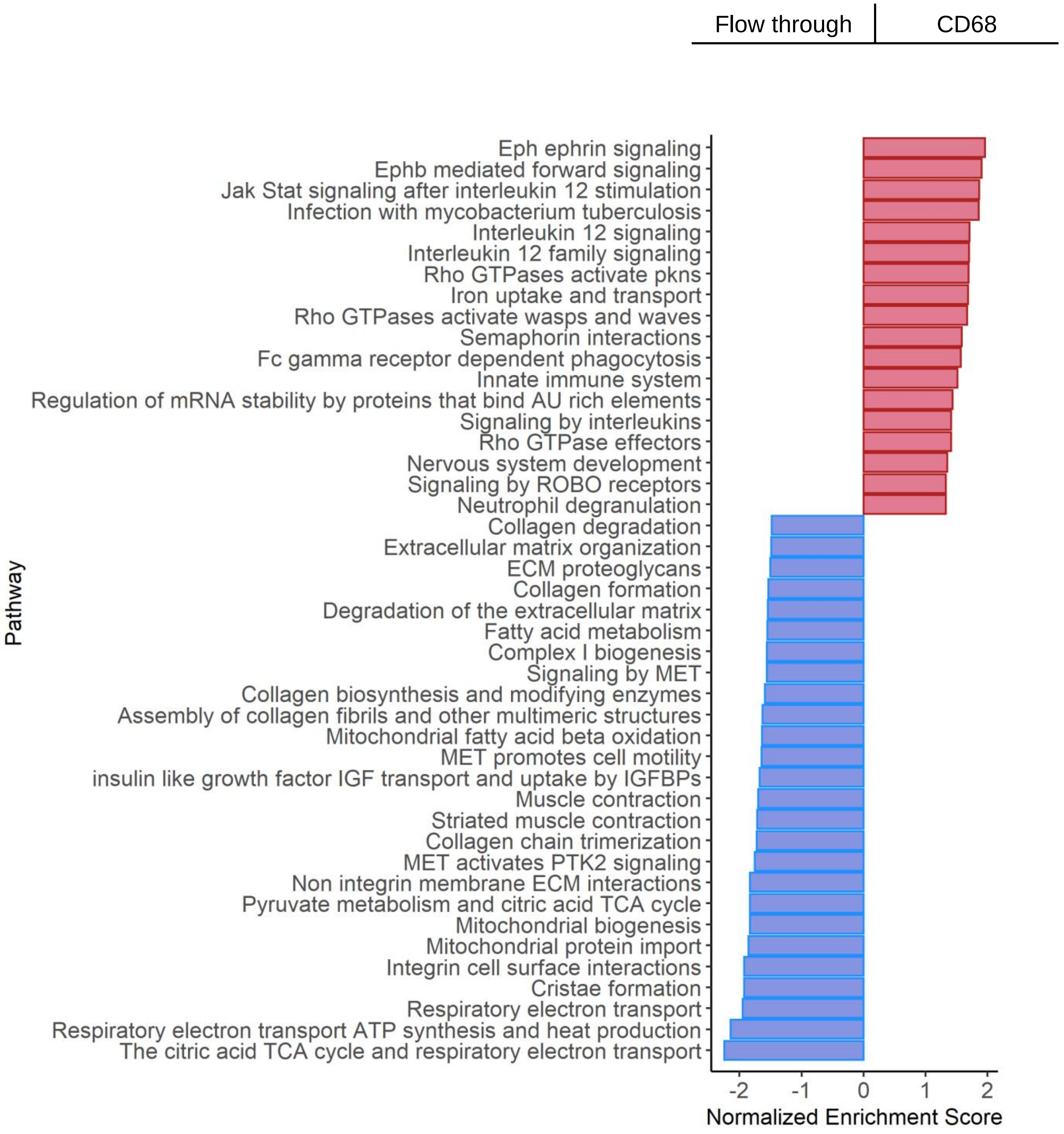


**Figure S3**. **Phagosome maturation and inflammatory signaling pathways are enriched in the CD68 fractions and mitochondrial metabolism pathways are enriched in the flow through fractions of rat cardiac infarcts**. All 1,671 of the proteins identified in the CD68 and flow through fractions (Figure 3D) were ranked by fold change of the LFQ value and searched by Gene Set Enrichment Analysis (GSEA) using the REACTOME annotation reference databases. The pathways are plotted by normalized enrichment score (NES) in which NES > 0 (red) indicate pathways enriched in the 'CD68' fraction and NES < 0 (blue) indicates pathways more abundant in the 'flow through' fraction.

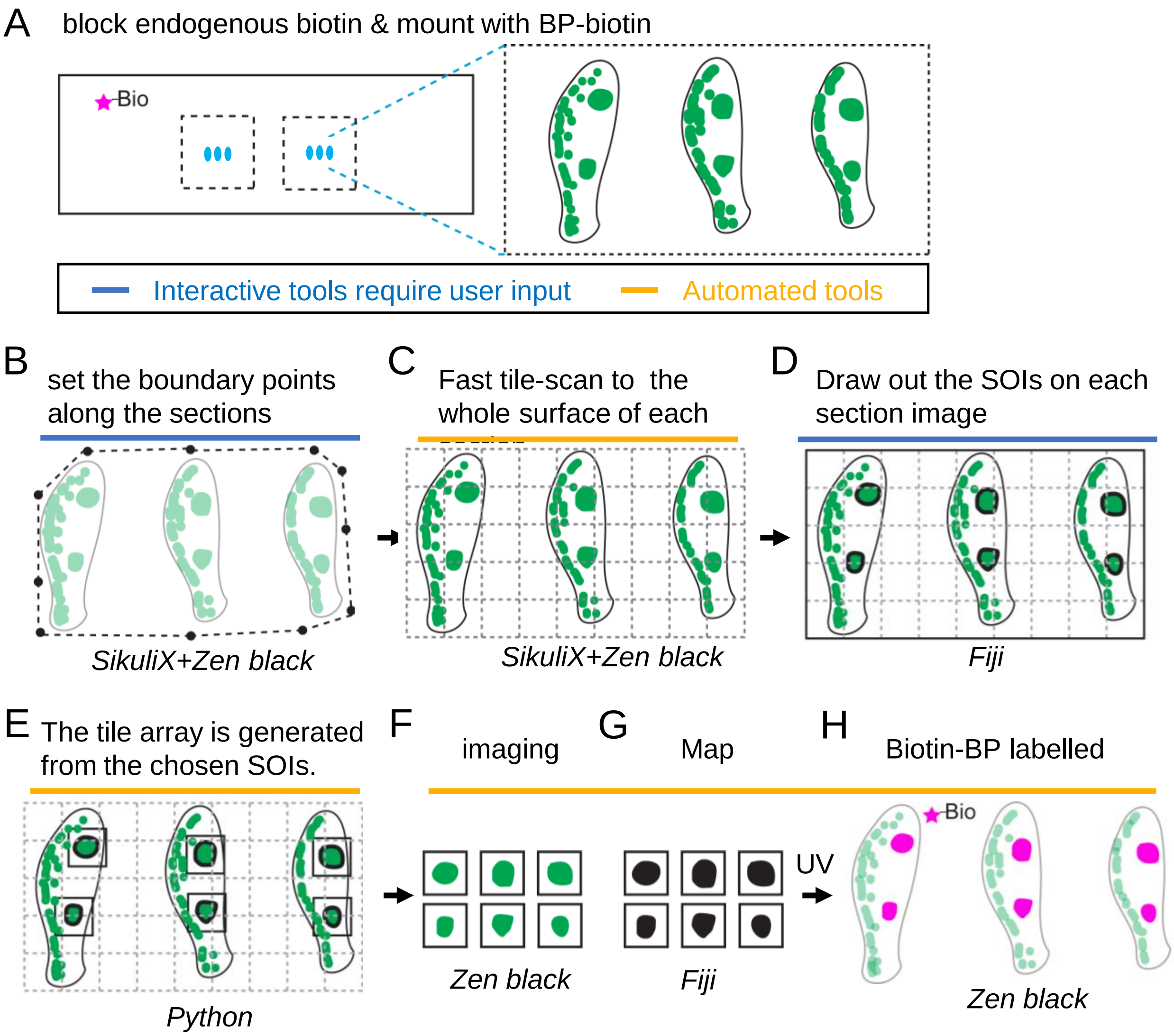


**Figure S4**. **Schematic of autoSTOMP 2.0 mediated targeting of discrete IgG4⁺ lesions across multiple esophageal biopsy sections.** 1 mm esophagus punch biopsies are isolated from a patient with active EoE. **A,** 6-8 sections per slide are stained for IgG4 (green), endogenous biotins are blocked and mounted with BP-biotin. A typical section contains 5 tiles at 340 μm x 340 μm per field of view using a 25x magnification objective lens. **B-H,** The SikuliX script allows the user to set the boundary points for all sections in Zen Black (**B**). **C,** The boundary points then direct low resolution tile scan in Zen Black. **D**, IgG4⁺ structures of interest (SOI) are identified by the user on each section in Fiji. **E,** A Python script maps the coordinates of each SOI to generate a tile array SOI-containing fields of view. **F-H,** A SikuliX script automates SOI imaging in Zen black (**F**, green), MAP file generation in Fiji (**G**, black) and BP-biotin crosslinking in Zen Black (**H**, magenta) as described in Figure 1.

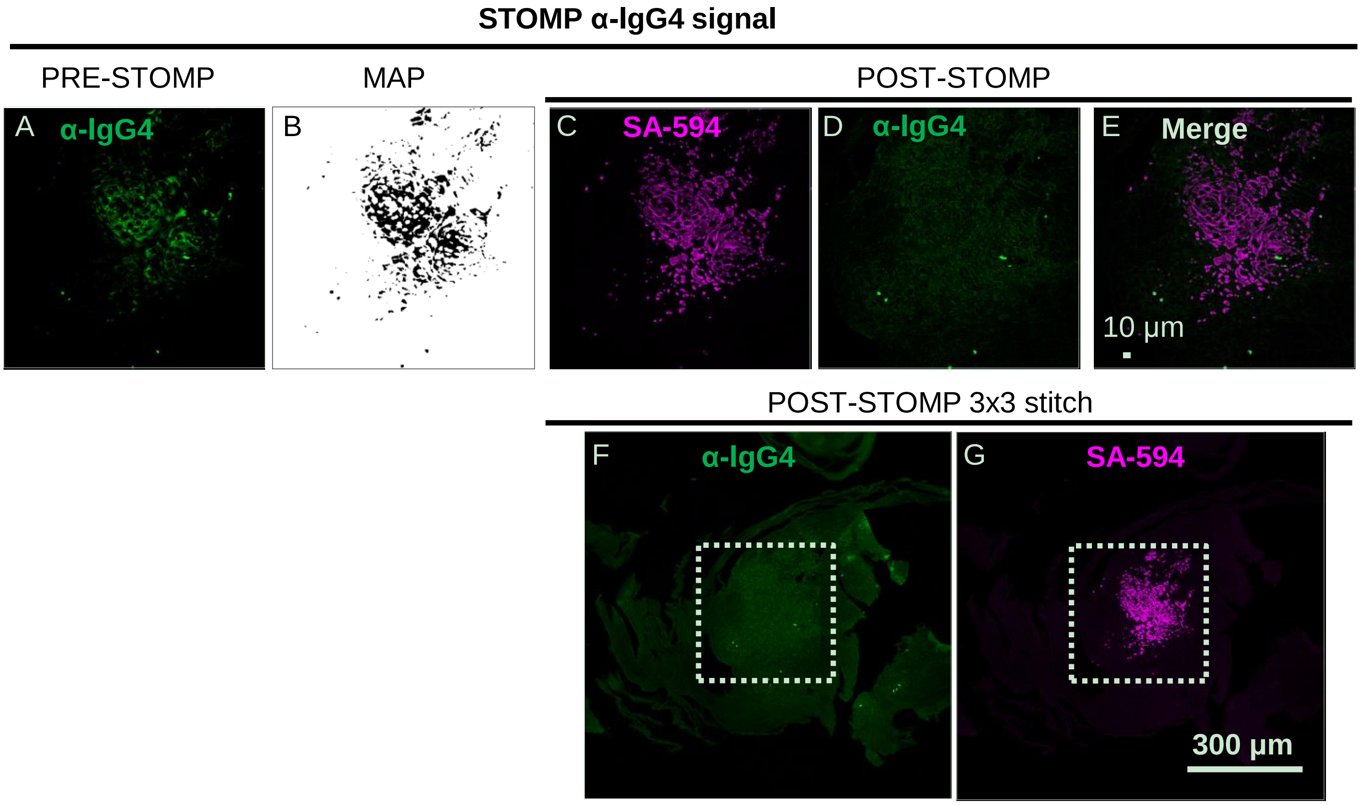


**Figure S5**. **AutoSTOMP2.0 selectively biotinylates IgG4+ regions of human EoE esophagus biopsies. A**, Representative image of an EoE esophagus biopsy section stained with an antibody specific to IgG4 (green). **B**, The MAP file generated in FIJI to identify the IgG4 signal in A. **C-E**, After UV-mediated BP-biotin tagging, the slide was washed to remove unconjugated BP-biotin, and biotinylated proteins were visualized using streptavidin-Alexafluor594 (**C**, SA-594, magenta). Samples were re-imaged for IgG4 signal (**D,** α-IgG4, green note photobleaching) and streptavidin 594 (**D**) co-localization **E**, merge. **F-G**, 3x3 tile array centered on the field of view targeted by the STOMP macro (white dotted line box) demonstrating lack of BP-biotin background in the targeted field of view relative to the surrounding regions. Representative of 3 independent experiments.

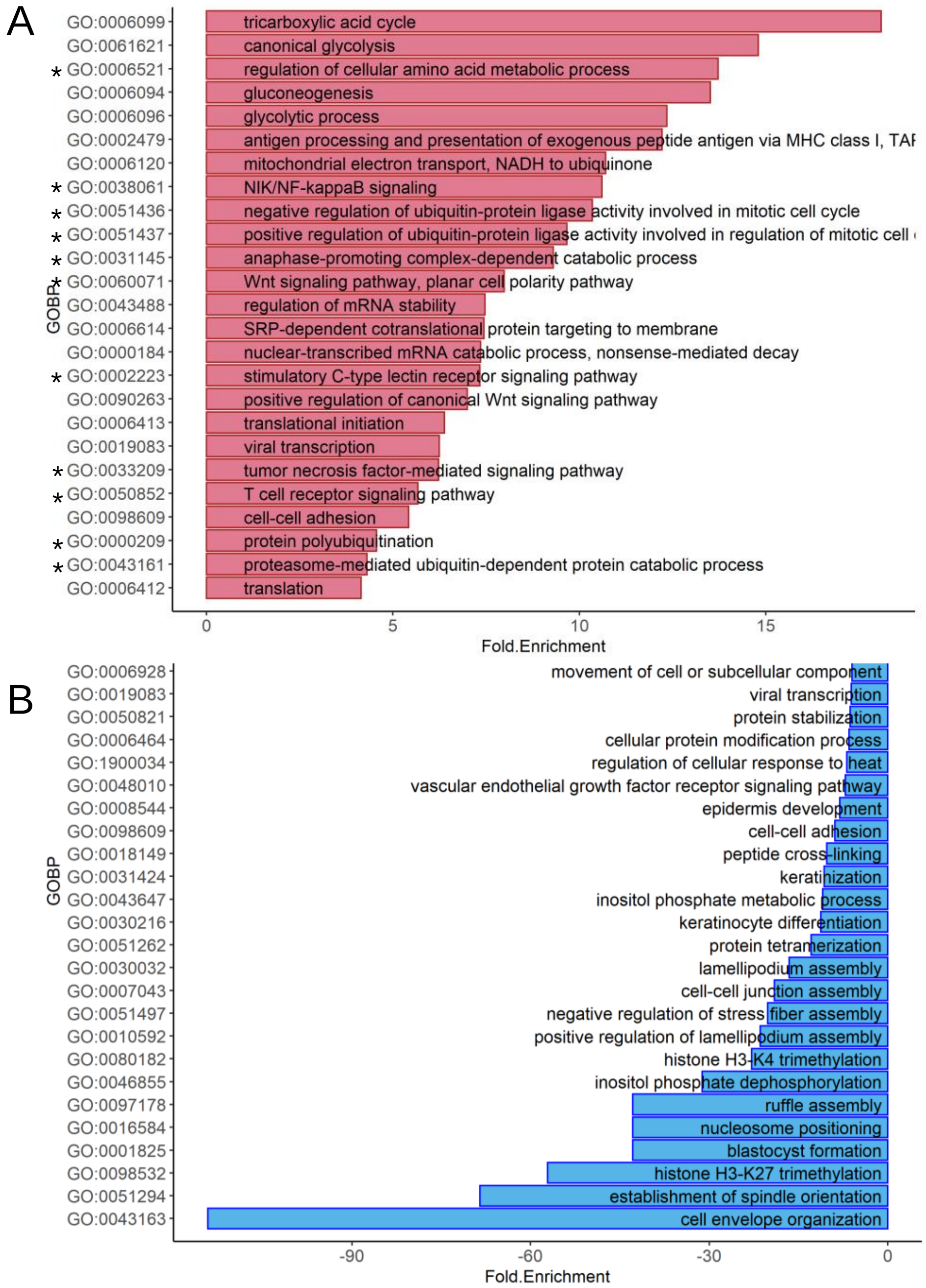


**Figure S6. Glycolytic metabolism and inflammatory pathways are enriched in IgG4⁺ regions of EoE biopsies and protein regulators of epithelial differentiation enriched in the flow through.** Proteins that were significantly differentially enriched (A, red) or depleted (B, blue) in the IgG4⁺ vs. flow-through (Figure 4) were searched against the Gene Ontology Biological Process (GOBP) terms using DAVID Bioinformatics Resource. The top 25 significantly enriched GO terms are displayed in a ranked by fold enrichment and false discovery rate (FDR). * indicates pathways defined by proteasome subunit representation.

#### Supporting Methods

##### Reagents and consumables

Chemical reagents and consumables were purchased from Thermo Fisher Scientific and used according to the manufacturer instructions unless specifically noted.

##### Creation of infarcts via LAD permanent ligation in the rat

Myocardial infarctions were performed in Eight-week-old male Sprague-Dawley rats weighing 275-300 g (Envigo RMS Division) after a one week acclimation period. On the day of the surgery, animals were anesthetized by intraperitoneal injection of ketamine (60-80 mg/kg) and xylazine (5-10 mg/kg). Animals were intubated and ventilated with oxygen and supplemental isoflurane (0.5-2%) as needed to maintain a proper depth of anesthesia. Bupivacaine was administered locally prior to the incision for pain management of the incision site (0.3 ml of a 0.25% solution). A left thoracotomy was performed, and the left anterior descending coronary artery (LAD) was ligated with 6-0 Monosof silk suture. The chest was closed using 3-0 resorbable suture to close the ribcage, 4-0 resorbable suture to close the muscle layer, and staples to close the skin (Medtronic, Minneapolis, MN). Buprenorphine (0.5-0.2 mg/kg) was administered subcutaneously immediately, with additional doses every 8-12 for 48-72 hours as required to manage postoperative pain. At 1 week following the initial ligation procedure, animals were anesthetized with 3.0% isoflurane in oxygen, intubated via tracheotomy, and ventilated with oxygen and 2.0-3.0% isoflurane. The chest was opened via a midline sternotomy. The heart was arrested via retrograde perfusion with cold BDM in PBS and removed for further processing. All experiments were approved by the University of Virginia Institutional Animal Care and Use Committee.

##### Rat infarct staining and histology

Scars were dissected from the arrested hearts, taking care to cut away as much of the viable muscle as possible. The scars were then frozen in a cryomold by submerging them in OCT and placing the cryomold in pre-chilled isopentane submerged in liquid nitrogen. Once frozen, scars that were not used immediately were stored at -80°C. The OCT blocks were then mounted onto a cryostat and cut into seven μm sections parallel to the epicardial surface. One slide from the mid-wall of each scar was fixed in acetone at -20°C for 15 minutes and stained with H&E to visualize the cell nuclei morphology. On an adjacent slide collagen deposition was confirmed by staining with picrosirius red to visualize the scar-muscle border in the sample. 6 to 8 slides from the mid-wall region of the scar were stained for the macrophage marker CD68. These sections were fixed in methanol at -30°C for 15 min. After washing in three washes in PBS, sections were then blocked for endogenous biotin using a blocking kit (Vector Laboratories). After washing three times with PBS, the slides were blocked in 2.0% BSA solution in TBS-0.1% Tween 20 (TBST) (Sigma-Aldrich). The CD68 antibody conjugated to Alexa Fluor 488 (clone:ED1, Bio-Rad Laboratories) was then applied to the tissue sections for one hour. The slides were washed three times with PBS then DI water and allowed to dry before being used for autoSTOMP. Slides that were not used immediately were stored at -30°C.

#### Eosinophilic esophagitis (EoE) patient biopsy preparation

The human study was approved by the University of Virginia Institutional Review Board (IRB). A written participant consent (IRB-HSR#19562) was filled by each participating patient at time of biopsy. Diagnostic criteria for EoE was a histological test demonstrating >15 plasma eosinophils per high power field (hpf) microscopy and dysphagia following the ACG clinical guideline[1]. Six 1mm biopsies were isolated using a standard endoscopic procedure.

The biopsy samples were rinsed in PBS, blotted on a paper towel and transferred to cryomolds in OCT. 3 to 4 biopsy samples were embedded in one OCT block and frozen in pre-chilled isopentane submerged in liquid nitrogen. nce frozen, scars that were not used immediately were stored at -80°C. The OCT blocks were then mounted onto a cryostat and cut into seven µm sections. Two sections from the OCT block were mounted onto each microscopic glass slide (6 to 8 tissue sections per slide, total).

The tissue sections were fixed in pre-chilled methanol for 15 minutes at -30°C then washed three times with PBS. Endogenous biotin was blocked using an avidin/biotin blocking kit (SP-2001, Vector Laboratories) per manufacturer's protocol. Sections were washed 3 more times PBS then blocked with 2% goat serum in TBST for 30 minutes. Sections were stained with an antibody specific to human $IgG_4$ (MRQ-44, mouse monoclonal, Cell Marque) in TBST at a dilution of 1:100 for 1 h in the dark. After 3 washes with TBST, the goat anti-mouse-AF488 secondary was diluted 1:300 in TBST and applied for 45 min in the dark. The slides were washed three times with TBST then DI water and allowed to dry before being used for autoSTOMP. Slides that were not used immediately were stored at -30°C.

#### Photo-activated biotin-BP cross-linking reaction

Biotin-dPEG-3-benzophenone (biotin-BP, 10267, Quanta BioDesign) was used as the UV activatable crosslinker to tag the proteins. A 0.5 M stock solution of biotin-BP was made in anhydrous DMSO (89139-666, VWR) and stored with desiccant in the dark at -30 ˚C, which is stable for 12 months. 1 mM biotin-BP mounting media was prepared freshly by diluting the biotin-BP stock solution in 50/50 (v/v) DMSO/water and used within 4 hours. Every cryosection was mounted with 12 µL biotin-BP mounting media, covered with cover glass (1.5 thickness, 18 mm diameter, #64-0714, Harvard Apparatus), and sealed with nail polish (i.e., Double Duty Base and Topcoat, Sally Hansen). The slides can be transferred to the microscope for autoSTOMP[2] after thermal equilibration for ~ 20 min. The mounted slide should not be stored longer than 24 h to limit potential for background reaction of biotin-BP.

Photo cross linking was performed on a Chameleon multiphoton light source (Coherent) coupled with an LSM880 confocal microscope (Carl Zeiss) in the Ewald Lab at the University of Virginia. A 25x oil immersion lens (LD LCI Plan-Apochromat 25x/0.81 mm Korr DIC M27) with immersion oil (518 F for 30°C, refractive index = 1.518, 444970-9000-000, Carl Zeiss) was used for imaging. Using the SikuliX[2] automation platform (version 2.0.0, http://sikulix.com/) tasks were integrated in Zen Black (Carl Zeiss), FIJI (FIJI Is Just ImageJ)[3], and Spyder (Python IDE, version 3.2.8, www.spyder-ide.org) using a user-friendly workflow with step-by-step guidance. The workflow was similar to the previous protocol[2]. Modifications for tissue sections included an additional module allowing the user to identify regions of interest and skip void regions within a tissue section as well as continuously sample multiple tissue sections per microscopic slide.

Please refer to the autoSTOMP2.0.docx protocol for tissue sections and biopsy samples and source scripts are available on GitHub (GitHub Inc.): https://github.com/boris2008/autoSTOMP_2.0.git

**Immunofluorescence (IF) validation of biotin tag cross-linking**

Following biotin-BP cross-linking, the coverslip was carefully removed by soaking in DI water in the dark at room temperature for at least 30 min. Sections were washed three times with 50/50 (v/v) DMSO/water and three times with Milli-Q water to rinse the biotin-BP mounting media off of the slide. The tissue sections were blocked in 1/20 (w/v) BSA/TBST for 30 min, and then stained with Alexa Fluor-594 Streptavidin (#016-580-084, Jackson ImmunoResearch Lab) at a 1:500 dilution in TBST for 45 min followed by three times washes with TBST. Before imaging, the tissue sections were mounted with mounting media containing DAPI (H-1000, Vector Laboratories) or stored at 4 $^{o}$C in the dark.

**Protein fractionation streptavidin precipitation**

Following biotin-BP cross-linking, the coverslip was carefully removed by soaking in DI water in the dark at room temperature for at least 30 min. Sections were washed three times with 50/50 (v/v) DMSO/water and three times with Milli-Q water. Excess water was aspirated, and the coverslip was stored at -30 ˚C while photo-crosslinking was performed on additional slides.

Two lysis protocols were developed specific to each tissue type. Lysis buffers were prepared freshly and used immediately for each batch of samples. Every 1 mg of rat cardiac sections were dissolved in 100 µL hydroxylamine lysis buffer[4] (1 M $NH_2OH$-HCl, 8 M urea, 0.2 M $K_2CO_3$, pH adjusted to 9.0) for 17h at 45 $^{o}$C with shaking at 800 rpm. The pH of the lysate was brought to 7.4. Insoluble debris was removed by centrifugation at 20,000 g for 15 min. EoE esophagus sections were dissolved in DTT/SDS buffer[5,6] (0.1 M Tris-HCl, pH=8.0, 0.1 M DTT, 4% SDS) at 1 mL/10 mg buffe/tissue weight. Samples were incubated at 99 $^{o}$C for 1 hour shaking at 600 rpm. After cooling to room temperature, the crude extract was clarified by centrifugation at 20000 x g at room temperature for 10 min.

For affinity precipitation, streptavidin (SA) magnetic bead (Pierce #88817, Thermo Fisher Scientific) were used. The lysate was diluted in TBST at a volume ratio of 1:9 and vortexed. Lysate was split into more than one 1.5 mL microtubes if the diluted lysate volume was larger than 1 mL. Every 15 to 20 µL of SA beads were used for a total tissue weight 1 to 2 mg (approximately 4 to 6 rat scar sections, or 16 to 28 EoE biopsy sections). The number of tissue sections used depended on the frequency of SOI in each section and the section size and thickness. The SA bead slurry was rinsed once with 1 mL TBST, then directly added to the diluted lysate and vortexed. Biotinylated proteins were precipitated for 1 hour at RT. The unbound lysate was collected as the ‘flow through’ fraction. The SA beads with bound SOI proteins (the ‘autoSTOMP’ fraction) were washed 7 times with TBS+0.1% SDS and 3 times with 100 mM ammonium bicarbonate (ambic) to deplete the nonspecifically bound proteins. The ‘autoSTOMP’ fraction on SA beads was resuspended in 100 µL 100 mM ambic buffer. The ‘flow through’ was pelleted by trichloro acetic acid (TCA) precipitation.

**LC-MS/MS analysis**

Both fractions were submitted to the University of Virginia Biomolecular Analysis Facility Core. In the core, the fractions were treated by on-bead or in-solution trypsin digestion and necessary desalting steps. The samples were reduced in 10 mM DTT at room temperature for 30 min followed by alkylation in 15 mM iodoacetamide in the dark at room temperature for 1 h. A 5 mM DDT solution was added to quench excess iodoacetamide. 1 µg of sequencing grade modified trypsin (V5111, Promega) was spiked into each sample. Digestion occured overnight (~12 h) at 37 ˚C. Formic acid was added at a final concentration of 1% (v/v) to stop the trypsinization. The sample was cleaned by SP3 protocol[4], dried. Salt is visible in the dried sample indicating further cleaning is required. In this case, C18 tips (#87784, Thermo Scientific™) was used for a second time desalting. The desalted peptide digest is reconstituted in 20 µL of 0.1% formic acid aqueous buffer. The samples were analyzed on a Thermo Electron Q Exactive HF-X mass spectrometer system with an Easy Spray ion source connected to a Thermo 75 µm x 15 cm C18 Easy Spray column. 10 µL (~ 50%) of the extract was injected and the peptides eluted from the column with a gradient at a flow rate of 0.3 µL/min at 40 °C for more than 1 hour. Solvent A (water + 0.1% formic acid) and solvent B (80% acetonitrile + 0.1% formic acid) were mixed in the instrument at various ratios (A+B = 100%) to create a gradient of eluation buffer. The gradient was applied in the order of 2% B for 1.5 min, 2-23% B for 51 min, 23-35% B for 10 min, 35-95% B for 1 min, and 95% B for 5 minutes. The nano-spray ion source was set at 1.9 kV, using a Top10 method with the MS scan at 120K resolution and HCD scans at 30K resolution. This mode of analysis produces approximately 25000 MS/MS spectra of ions ranging in abundance over several orders of magnitude.

**MS Data Analysis**

The Thermo MS "RAW" file was analyzed using the andromeda engine in MaxQuant (version 1.6.14.0, Max Planck Institute of Biochemistry)[7]. Common contaminants (http://www.coxdocs.org/doku.php?id=maxquant:start_downloads.htm) were searched to be excluded from our result. Peptide FDR and protein FDR were both set to 1%. Protein abundance was reported as label-free quantification (LFQ)[8] and normalized between the fraction 'autoSTOMP' and 'flow through' (three replicates). The default MaxQuant global parameters and group-specific parameters were used. For rat scar samples, the raw data were searched against the Rattus norvegicus Uniprot Fasta file (up000002494). In addition, counting of the $NH_2OH$ and trypsin enzymatic cleavage sites[4] (2 maximum missing cleavage sites allowed) and post-translational modifications (PTMs) including fixed PTMs (cysteine carbamidomethylation) and dynamic PTMs (methionine oxidation, hydroxylysine, and hydroxyproline) were used. For EoE samples, the raw data were searched against the Homo sapiens Uniprot Fasta file (up000005640) with counting the trypsin cleavage sites and PTMs on methionine and cysteine.

The LFQ data in the MaxQuant output, "proteinGroups.txt" was analyzed in Perseus (version 1.6.14.0, Max Planck Institute of Biochemistry)[9] following the software recommended protocol[9]. Similarity between the three replicates of the 'autoSTOMP' and 'flow-through' fractions was determined by t-distributed stochastic neighborhood embedding (t-SNE)[10]. The differential expression of each protein was analyzed with Student's t-test with significance defined by permutation-based FDR (< 0.05).

Data were visualized with R package “ggplot2”, ”ggrepel” and “heatmap.2” using R (www.r-project.org) or GraphPad Prism (version 8.2.1).

**Immunofluorescence (IF) validation of autoSTOMP results**

To validate co-localization of candidate proteins with CD68 or IgG4 in rat tissue section or human biopsy, methanol fixed sections were washed three times with PBS and permeabilized in 0.1% Triton-X100 in PBS at room temperature for 15 min. Samples were blocked in 5% BSA in TBST (TBS+0.1% tween20) at room temperature for 30 minutes. The primary antibody was diluted in TBST and incubated at room temperature for 1.5 hours, washed three times in TBST and stained with secondary antibodies for 1 hour at 1:500 dilution. Samples were washed three times in TBST and mounted in DAPI-containing mounting media (Vector Laboratories #H-1200-10).

Table 1. The antibodies used on rat scar tissues.

| Sequential applied | Lyz2 & Cd68 | Lamp1 & Cd68 | C1qc & Cd68 |
|---|---|---|---|
| 1 | Rbt-α-Lyz2 (Invitrogen#PA5-114441) | Rt-α-Lamp1 (BD Pharmingen #553792) | Rbt-α-C1qc (Novus Biologicals#NBP2-30024) |
| 2 | Gt-α-Rbt-AF594 (Invitrogen# A11037) | Gt-α-Rt-AF594 (Invitrogen# A11007) | Gt-α-Rbt-AF594 (Invitrogen# A11037) |
| 3 | M-α-CD68-AF488 (#MCA341a488, bioRad) | M-α-CD68-AF488 (bioRad,#MCA341a488) | M-α-CD68-AF488 (#MCA341a488, bioRad) |

### host: rabbit (Rbt), rat (Rt), goat (Gt), mouse (M).

Table 2. The antibodies used on Eoe samples.

| Sequential applied | Prg2 & IgG4 | Alox15 & IgG4 | IL-1RA & IgG4 |
|---|---|---|---|
| 1 | Rbt-α-Prg2 (proteintech #10766-1-AP) | Rbt-α-Alox15 (Sigma life science#HPA013859) | Rbt-α-IL-1RA (proteintech#10844-1-AP) |
| 2 | Gt-α-Rbt-AF594 (Invitrogen # A11037) | Gt-α-Rbt-AF594 (Invitrogen # A11037) | Gt-α-Rbt-AF594 (Invitrogen # A11037) |
| 3 | M-α-IgG4 (Cell Marque # MRQ-44) | M-α-IgG4 (Cell Marque # MRQ-44) | M-α-IgG4 (Cell Marque # MRQ-44) |
| 4 | Gt-α-M-AF488 (Abcam #ab150117) | Gt-α-M-AF488 (Abcam #ab150117) | Gt-α-M-AF488 (Abcam #ab150117) |

* human biopsy section is blocked with human Fc blocker (BD #564220) before staining.

* host: rabbit (Rbt), rat (Rt), goat (Gt), mouse (M).
